# A Whole-Brain Dynamical Framework Linking Resting-State Activity to TMS-Evoked Responses

**DOI:** 10.64898/2026.07.10.737755

**Authors:** Andrea Veronese, Davide Momi, Simone Sarasso, Maurizio Corbetta, Michele Allegra, Samir Suweis

## Abstract

A major challenge in systems neuroscience is understanding how external perturbations interact with ongoing brain activity. Transcranial magnetic stimulation (TMS), increasingly used in both basic and clinical neuroscience and often combined with electroencephalography (EEG), provides a unique opportunity to probe this interaction. However, how intrinsic dynamics constrain the propagation of TMS-evoked activity remains poorly understood. In particular, effective connectivity (EC)—capturing directed, state-dependent interactions between brain regions—is thought to critically shape perturbational spread, yet remains difficult to estimate at the whole-brain EEG level. Here we introduce an analytically tractable, generative whole-brain model that links spontaneous EEG activity to cortical responses under perturbation. By deriving a closed-form expression for the model’s cross-spectral density, we directly fit empirical resting-state EEG spectra and infer biophysically interpretable local dynamical parameters without time-domain simulations. We then estimate stimulation-site-specific EC using only a small fraction of the TMS–EEG trials. The resulting model accurately predicts the spatiotemporal structure of TMS-evoked potentials (TEPs) in unseen trials. Moreover, even without subject-specific refitting, group-level EC templates capture canonical site-specific propagation motifs underlying single-subject early TMS responses. Together, our results establish an analytical framework for individualized whole-brain modeling of TMS-EEG with potential applicability to model-based neuromodulation.

## Introduction

Transcranial magnetic stimulation (TMS) combined with electroencephalography (EEG) provides a unique window into how local cortical perturbations propagate through large-scale brain networks. A single magnetic pulse induces a transient electric field that activates cortical neurons, generating a cascade of activity measurable with millisecond temporal resolution [1]. The resulting TMS–evoked potentials (TEPs) reflect both early stimulus-locked components, associated with the direct activation of the targeted region, and later components, linked to recurrent processing and long-range network interactions. Experimental studies have leveraged TMS–EEG to explore diverse cognitive and clinical phenomena [2, 3, 4, 5], establishing it as a powerful perturbational approach for mapping causal network interactions in the human brain [6, 7]. Observing distributed responses to stimulation has uncovered network features associated with cognitive performance [8], markers of consciousness [9, 10], and variations in long-distance cortical interactions [11]. Clinically, TMS-EEG is used to investigate the effects of repetitive TMS in treating major depression [12], schizophrenia [13], and disorders of consciousness [14].

Despite this widespread application, the mechanisms supporting the observed whole-brain TMS responses are still only partially understood. Empirically, TMS responses depend strongly on the brain’s ongoing state [6, 15, 16], suggesting that the same intrinsic network dynamics that govern the resting state also constrain the impact of a perturbation. Understanding this relationship is critical for addressing the limited predictability of clinical TMS interventions [17] and optimizing individualized stimulation targets [18]. In this context, whole-brain computational models represent a promising tool [19]. By combining local neural dynamics with large-scale connectivity, these models attempt to capture the mechanisms underlying distributed brain activity, both at rest and under the effect of external stimuli and perturbations [20, 21, 22, 23, 24], potentially bridging spontaneous and evoked brain activity. Among the relevant parameters, whole-brain models often include a principled estimation of effective connectivity (EC), capturing directed and state-dependent interactions between brain regions. In TMS–EEG, EC is expected to play a crucial role in determining how perturbations propagate through cortical networks and how stimulation of a given region recruits distant areas.

Currently, no model simultaneously encompasses whole-brain resting-state EEG activity and TEPs. In fact, classical dynamic causal models [25, 26] infer effective connectivity (EC) modulated by external conditions (such as rest vs. stimulation). Still, they are limited to 10-12 nodes due to model complexity and, hence, are unsuitable at the whole brain level. The Virtual Brain allows fitting dynamical models on whole-brain EEG activity but assumes a fixed inter-areal coupling given by the anatomical connectivity [27], which limits the accuracy in reproducing the observed resting-state spectra and TEPs. Recent works [28, 29] fit a dynamic model directly to TEPs, achieving high fidelity but without ensuring that the model can simultaneously account for the underlying spontaneous dynamics. A common element of all these models is that they are based on biophysically grounded dynamical models of local neural masses, which renders inversion of the whole-brain model analytically intractable, requiring sophisticated inference techniques. The fast timescales of electrophysiological signals, the strong impact of volume conduction, and the high dimensionality of source-resolved data make model inversion highly sensitive to parameter degeneracy and overfitting. As a consequence, inferred EC patterns may become poorly constrained or physiologically implausible. Developing analytically tractable whole-brain models that allow stable and interpretable estimation of effective connectivity from EEG data is, therefore, a critical step toward understanding how intrinsic brain dynamics shape responses to external perturbations.

To bridge this gap, we present a generative, tractable modeling framework that reliably reproduces the resting state spectral properties of the human brain while maintaining the capacity to simulate evoked responses. Our approach combines the Hopf model [30] with a two-step training protocol (Figure 1). In the first step, we fit the model to each subject’s resting-state EEG. Due to the simplicity of the model, which represents each brain region as a noisy Stuart-Landau oscillator, we obtain an analytical solution for the cross-spectral density, enabling the precise fitting of the model to the resting-state covariance matrix and spectral power of each subject. The fit yields subject-specific values for the local parameters of the model, and a subject-specific effective connectivity (EC). In the second step, we keep the local model parameters fixed at their resting-state values, and use a small fraction (10%) of TMS–EEG trials to re-fit the EC, constraining it to remain close to the resting-state baseline. This second fit yields a subject- and stimulation-site-specific EC required to predict the remaining 90% of unseen TEP data. The first step, which distinguishes our approach from previous work, provides several advantages. First, it is the key to obtaining a model that jointly captures spontaneous and stimulation-evoked activity. This model allows conceptualizing TEPs as resulting from a modulation of the unperturbed whole-brain dynamics, and demonstrates how the combination of resting-state dynamics and a few stimulation trials enables robust simulations of whole-brain responses to TMS Second, it enhances the accuracy of the model: when fitting the model directly on the TEP without accounting for the resting-state baseline (e.g., using anatomical connectivity to constrain the EC), the model quality is consistently degraded.

**Fig. 1.**
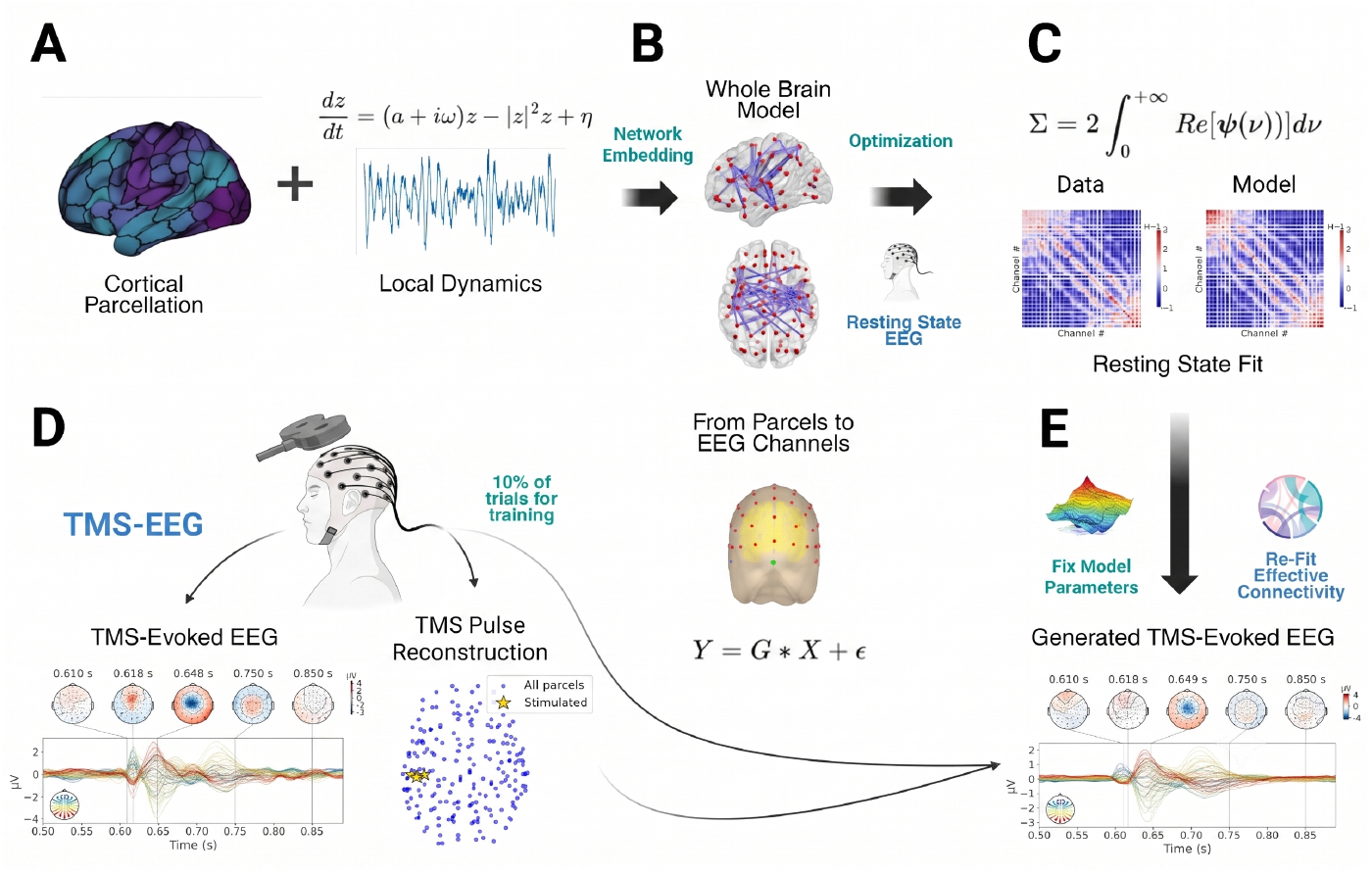
Methodological workflow. Subject-specific connectome-based neurophysiological modelling of resting state EEG and generation of TMS–EEG TEPs. **(A)** The cortical surface is parcellated according to the Schaefer-200 atlas, with each parcel modelled as a network node governed by Hopf local dynamics. **(B)** The whole-brain model is constructed by embedding these nodes into a network defined by structural connectivity (SC) and individualized connectivity gains. This stage includes the derivation of the source-to-sensor mapping to project simulated source activity into the EEG channel space. **(C)** Model parameters are optimized by fitting the analytical covariance derived from the linearized system to the empirical resting state EEG covariance matrix, establishing the intrinsic dynamical scaffold. **(D)** The TMS pulse is reconstructed in source space by weighting parcels according to electric field magnitudes. A small subset (10%) of empirical TEP trials is selected for connectivity refinement. **(E)** Keeping fixed all local and global parameters derived from the resting, the effective connectivity obtained at rest is re-optimized using the 10% TEP training subset. The resulting generative model is driven by the reconstructed pulse to predict the full spatiotemporal evolution of the TEP for the remaining 90% of unseen empirical trials.

Our model is demonstrated on a cohort of 11 subjects undergoing TMS split between Pre-frontal and Motor stimulation sites [31]. Beyond individualized fits, we demonstrate that the inferred site-specific *EC*_TMS_ can be aggregated to define group-level “propagation motifs”. By combining (i) a subject’s unique resting-state parameters with (ii) a data-driven group-average *EC*_TMS_ template and (iii) biophysically informed TMS input (SimNIBS [32]), it is possible to capture a meaningful component of the early phase of TMS-evoked responses at the single-subject level without any individual connectivity refitting (thus avoiding step 2). This group-average generalizability reveals a shared, reproducible architecture in early cortical reactivity, while simultaneously exposing the clear boundaries where individual, state-dependent variations begin to dominate later components of the signal.

## Results

### A tractable whole brain model for EEG

The dataset used consists of resting-state EEG recordings pooled from two separate experimental cohorts. The first cohort contributed resting-state data prior to a Prefrontal stimulation protocol (*n* = 6). The second cohort contributed resting-state data prior to a dedicated Motor stimulation protocol (*n* = 6). Following the exclusion of one motor baseline recording due to excessive artifacts, a total of 11 usable resting-state datasets were retained for final analysis. We modelled each cortical parcel as a Hopf oscillator near a supercritical bifurcation and coupled the parcels via a structural connectivity matrix augmented by delays. By linearizing the coupled system around its global fixed point (see Supplementary Note 1), we derived a frequency-dependent representation that allows direct computation of the resting state covariance matrix and spectral properties in sensor space. Unlike previous approaches that fit parameters directly to evoked responses, we first performed a “blind” fit to the resting state EEG (see Supplementary Note 2) for each subject to estimate a comprehensive set of parameters: node-specific excitabilities (*a*_*i*_) and intrinsic frequencies 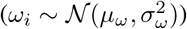, the global connectivity scaling (*g*), the standard deviation of the input noise (*σ*_*in*_), the homogeneous conduction velocity (*v*_*d*_) governing signal propagation delays (*m*_*ij*_), and a baseline connection gain matrix (*W*_*ij*_). Across the 11 available resting-state recordings, the model captured the resting-state EEG covariance structure with high fidelity (group mean *R*^2^ = 74% ± 9%, range [0.57, 0.88]). These results represent performance on a held-out test dataset derived via block sequential splitting, where the model was fitted using covariance matrices from an initial 20% of the samples and evaluated against the remaining 80% of the resting-state EEG timeseries. The model accurately reproduced the fronto-central and parieto-occipital motifs typical of spontaneous neural networks (Fig. 2A). Interestingly, the fitted parameters indicate supercritical local dynamics (*a*_*i*_ *>* 0, see Fig. 2B), establishing a self-sustained oscillation regime. This suggests that the resting-state alpha rhythms are generated by active limit-cycle oscillations rather than being externally driven by noise (as in the subcritical regime observed in fMRI). Unlike noise-driven models that rely on external stochastic input to maintain activity, this limit-cycle regime provides a stable dynamical scaffold that can be systematically perturbed by TMS, ensuring that the resulting TEPs are shaped by the network’s inherent resonance properties. Node-specific intrinsic frequencies were found predominantly in the alpha range (*μ*_*ω*_ ≈ 10 Hz), and conduction velocities (*v*_*d*_ ≈ 6.1-6.5 m/s) were consistent with myelinated white matter propagation. The effective connectivity matrix (see Supplementary Note 3) was characterized by dominant within-hemisphere coupling and a modular block structure aligning with the coarse anatomical segregation of major cortical systems (Fig. 2C). Notably, these parameters were estimated solely from the cross-spectral density of the resting data, providing a purely intrinsic scaffold for the subsequent perturbational modeling. The decoupled and non-physiological spatial patterns in (B) and (C) demonstrate how unconstrained mathematical optimization can achieve high temporal fitting accuracy by forcing structurally ungrounded and biologically meaningless network configurations. When trying to fit the model on phase-scrambled resting-state EEG data, where the empirical power spectrum is preserved but the correlation structure is destroyed (Supplementary Note 4), the optimization procedure failed to converge. This indicates that one cannot force the network of coupled oscillators to reproduce any multivariate time series by simply adjusting the model’s parameters, thus strengthening the parameters’ interpretability.

**Fig. 2.**
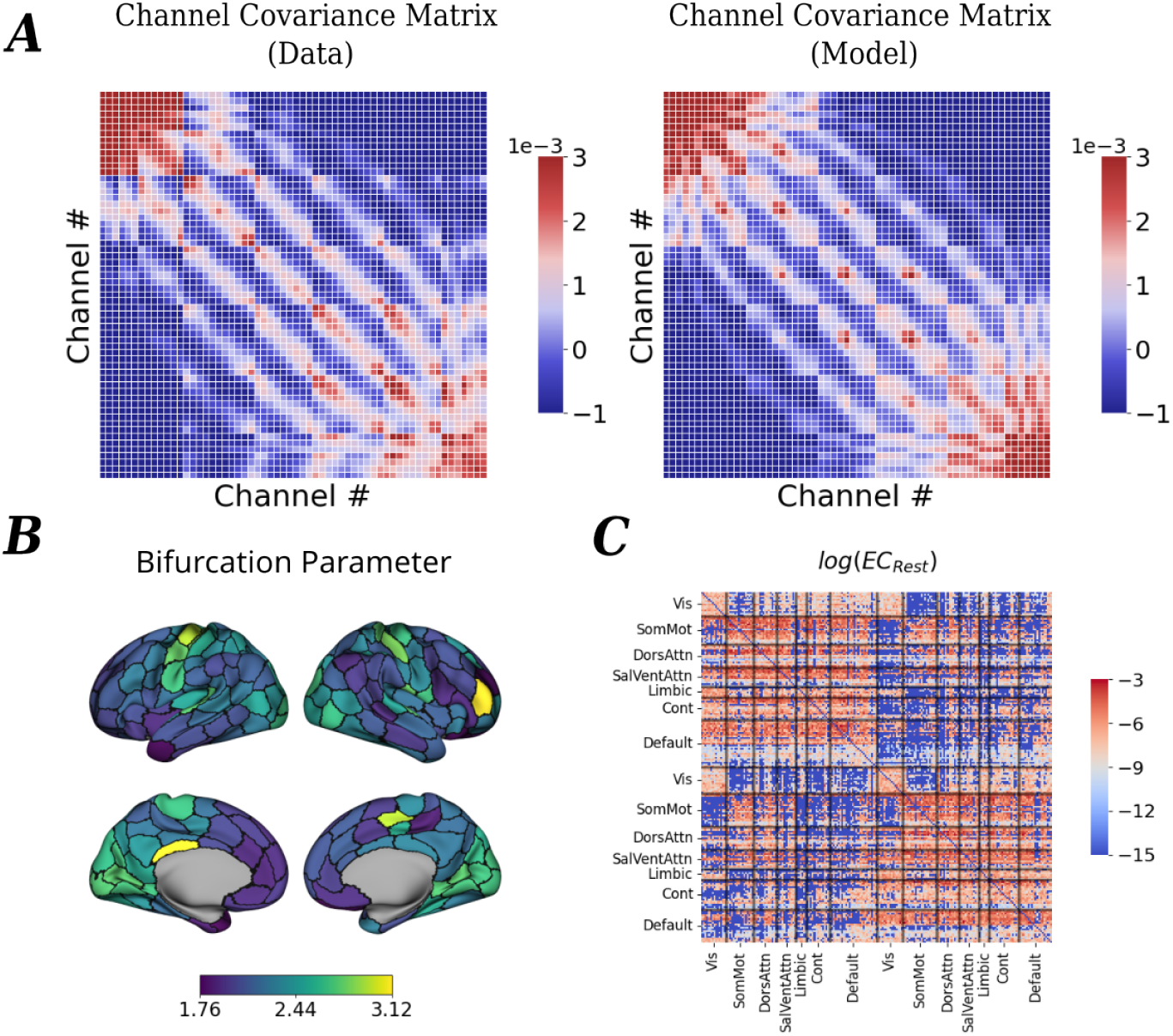
Hopf Whole Brain model resting state fit. (A) Empirical (left) and model (right) covariance matrices for a representative subject; the model captures the intrinsic spectral fingerprint. (B) Spatial distribution of the bifurcation parameter *a* on the Schaefer-200 parcellation, reflecting local neural population excitability. (C) The inferred effective connectivity (log *EC*_*Rest*_) exhibits a biologically plausible structure.

### Simulating TMS-evoked responses across stimulation sites

Having established that the Hopf model captures the resting state spectral structure, we next sought to determine if it can also capture the brain’s response to TMS. To do so, we fixed all parameters obtained during the resting state fit (*a*_*i*_, *μ*_*ω*_, *σ*_*ω*_, *g, σ*_*in*_, and *v*_*d*_) and refitted only the individual connection gains (*W*_*ij*_) using a randomly selected small subset (10%) of the TMS-EEG trials for each stimulation site, Prefrontal and Motor (Supplementary Note 5). This training sample size was selected based on a sensitivity analysis showing that model generalization performance reaches a stable plateau by this threshold (Fig. S1). Importantly, the connectivity refitting was constrained to remain near the resting state backbone to ensure biological plausibility and avoid overfitting (Fig. S2). However, we emphasize that the stimulation-induced effective connectivity was explicitly estimated from TMS data, and not derived purely from the resting state; it represents the necessary network reconfiguration required to propagate the external pulse. Using the parameters derived from the resting state and the refined EC, the model was tested on the remaining 90% of unseen TMS trials, evaluating goodness-of-fit metrics (*ρ* and *R*^2^) within a post-stimulus window spanning 0-250 ms relative to TMS onset. Crucially, for the Motor site, this fitting was restricted exclusively to low-MEP trials to isolate central cortical reactivity from peripheral somatosensory feedback.

The model successfully predicted the temporal profile and the topography of empirical TEPs for both prefrontal and motor stimulation (Fig. 3). For motor stimulation, simulated responses reproduced the triphasic waveform (N15, P30, and N45), matching empirical component latencies (simulated N15: 17.1 ± 7.7 ms; empirical 14.1 ± 3.7 ms). The model achieved robust fit performance across subjects (mean *R*^2^ = 0.77 ± 0.08, range [0.65, 0.87]), and spatial correlations between simulated and empirical spectral power were high across all bands (all *r >* 0.82, *p <* 0.001). Time-frequency analysis captured the canonical sequence of broadband suppression and subsequent frequency-specific rebounds (Supplementary Note 6). For prefrontal stimulation, the model again reproduced the triphasic waveform and obtained strong spatial correlations across frequency bands (all *r* > 0.82, *p* < 0.001), correctly capturing the heightened theta synchronization characteristic of frontal networks. However, the overall predictive accuracy for prefrontal targets (mean *R*^2^ = 0.50 ± 0.17, range [0.24, 0.78]) was lower than that for motor stimulation. This performance gap indicates that while the resting-state scaffold successfully predicts basic propagation motifs, prefrontal stimulation may engage more complex, state-dependent, or non-linear recurrent dynamics that are less fully captured by static connectivity adjustments.

**Fig. 3.**
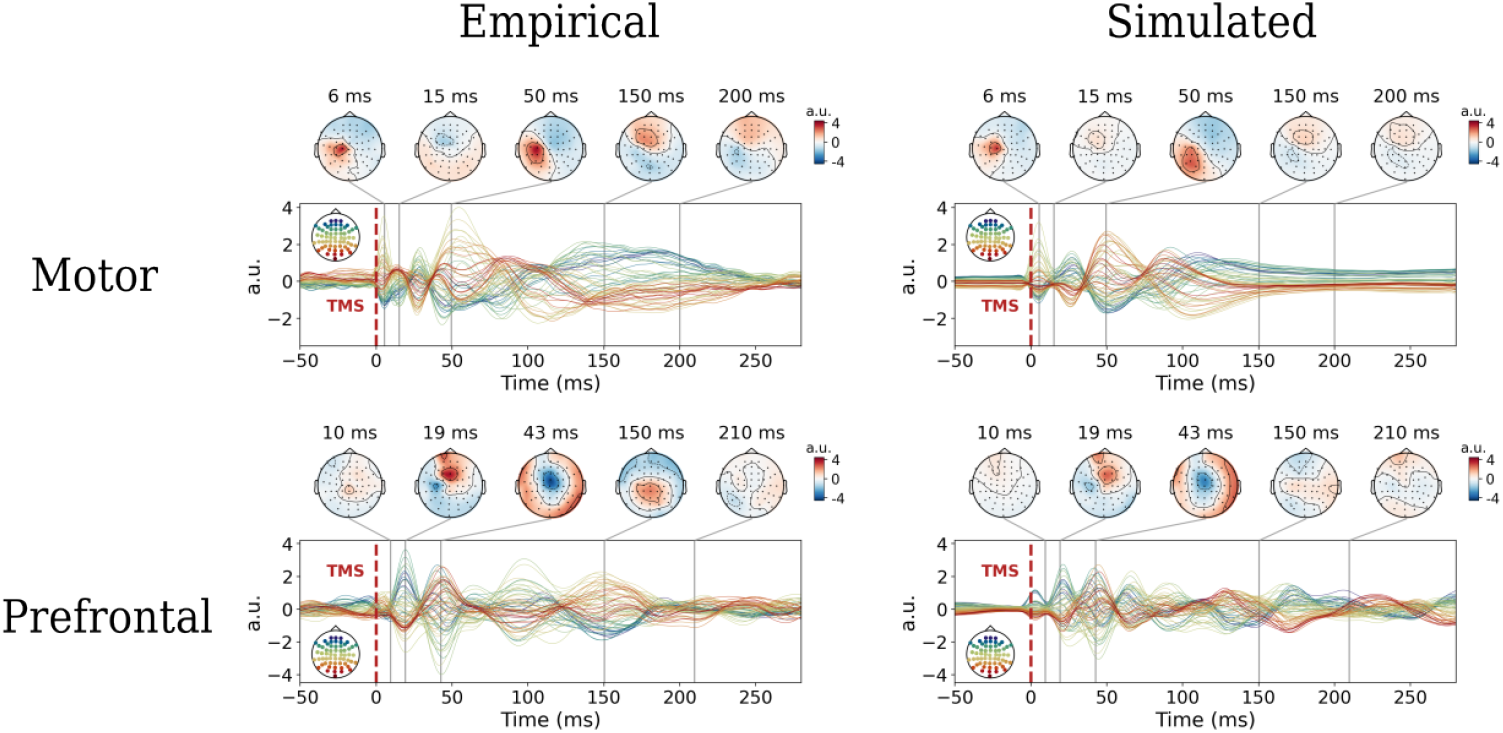
Site-specific TMS model simulations. Comparison of empirical (left column) and model (right column) responses in a representative subject (Subject 1). Temporal TEP correspondence for Motor (top row; *ρ* = 0.88 and *R*^2^ = 0.65) and Prefrontal (bottom row; *ρ* = 0.76 and *R*^2^ = 0.57) stimulation.

To evaluate the spatial and topological specificity of this fitting framework, we performed a cross-site generalization test using empirical data from two additional stimulation sites, Premotor and Parietal (not included in the main analysis because explicit SimNibs structural pulse reconstructions were unavailable). We tested whether the optimization framework initialized with the Motor pulse configuration could generalize to these alternative datasets. The model demonstrated a gradient of spatial sensitivity: it generalized remarkably well to the Pre-motor site, yielding stable predictive accuracy (mean *R*^2^ = 0.57 ± 0.11, range [0.34, ± 0.75]). Conversely, when evaluated on the anatomically and structurally distinct Parietal site, the model failed completely, yielding negative *R*^2^ values across subjects. This clear divergence indicates that while the model exhibits a degree of spatial tolerance for neighboring cortical regions within shared macro-scale networks (i.e., motor and premotor systems), it retains strict topological specificity and cannot fit unaligned, distant target dynamics.

Individual results for all subjects and stimulation sites are provided in Supplementary Figures S3–S13.

### Importance of spontaneous activity prior and of personalized EC

We investigated whether the successful simulation of the TEP is fundamentally dependent on the underlying resting-state activity prior. To test this, we evaluated the performance of our fully optimized model in predicting the TEP against three distinct control conditions (evaluated using Bonferroni-corrected t-tests).

First, to establish a baseline structural null condition (SC), we bypassed the resting-state effective connectivity entirely, utilizing only the empirical structural connectivity (SC) alongside local parameters averaged across subjects. Compared to the personalized whole brain model (WBM), where resting state parameters are inferred from each individual, the SC condition exhibited a severe drop in performance across both regions. This reduction was statistically significant for *ρ* in both Prefrontal (*p <* 0.05) and Motor (*p <* 0.01), as well as for *R*^2^ in Motor (*p <* 0.01), whereas the decrease in Prefrontal *R*^2^ did not reach statistical significance. Overall, these results highlight that structural connectivity alone, without the functional weighting provided by the *EC*_*rest*_ prior, is insufficient to fully capture the complex spatiotemporal dynamics of the evoked response.

Next, we assessed the subject-specificity of this resting prior by performing a parameter-swapping surrogate analysis (“Rest Params”). Here, we paired a subject’s true TMS-refined EC with the resting-state parameters derived from a different individual. We found no statistically significant difference in performance between the personalised WBM and the Rest Params condition (n.s.) for either metric or stimulation site (see Figure 4). This lack of sensitivity to individual resting-state parameter configurations is consistent with the fact that the underlying empirical resting-state covariance structure is highly conserved across the cohort, exhibiting remarkably high similarity between different subjects (see Figure S14). Because different individuals share a highly preserved resting state functional covariance matrix, swapping parameters still provides a structurally valid and plausible EC prior, allowing the optimization procedure to successfully fit the data.

**Fig. 4.**
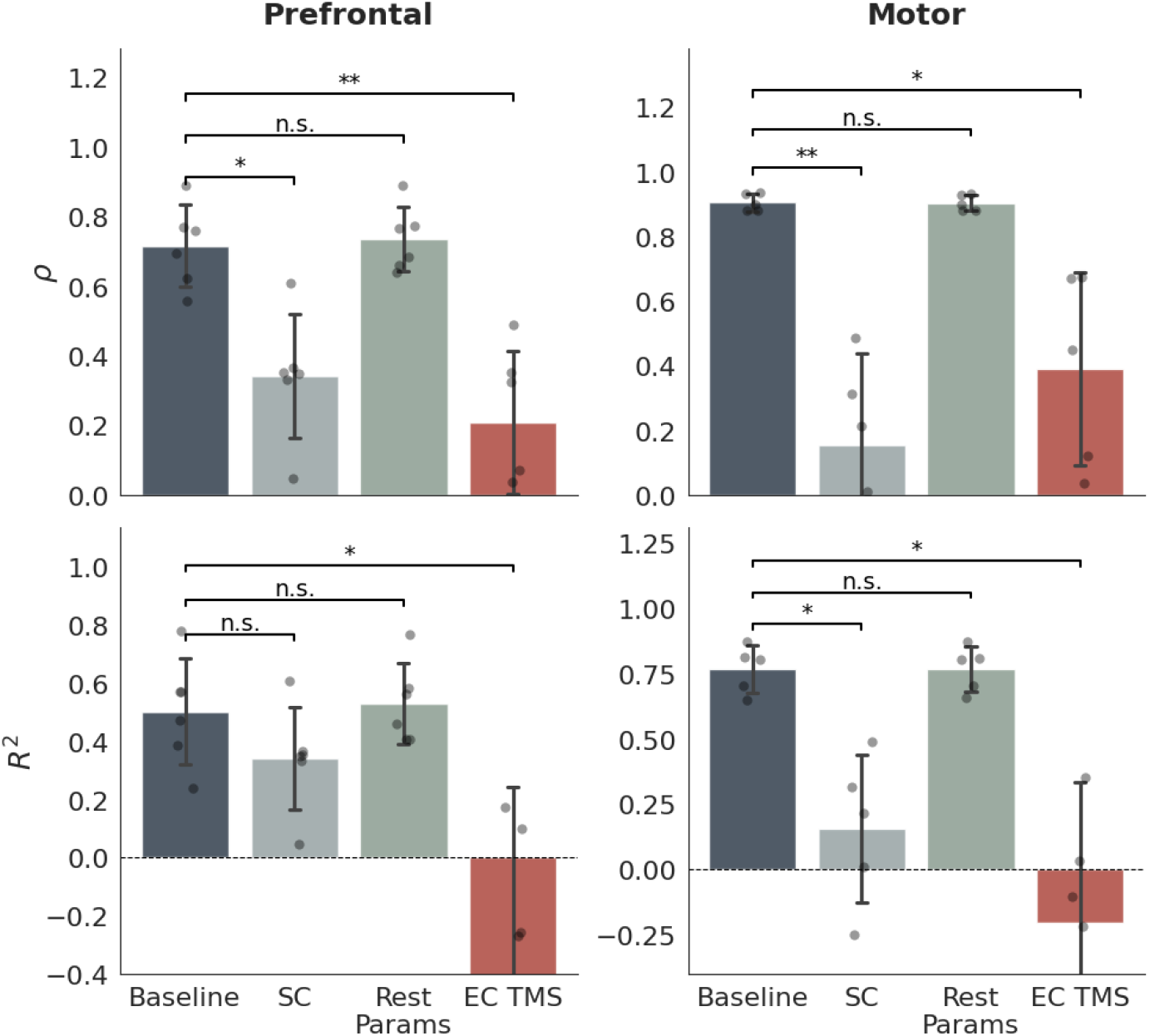
Evaluation of null conditions. Group-level goodness-of-fit metrics comparing the fully optimized personalised WBM against three control conditions: SC (structural connectivity only, paired with group-average local parameters), Rest Params (subject-specific, TMS-refined effective connectivity (EC) paired with donor resting-state parameters), and EC TMS (subject-specific resting WBM paired with donor TMS-refined EC). (Top) Pearson correlation (*ρ*) and (Bottom) Coefficient of determination (*R*^2^) between empirical and simulated TMS-evoked responses for the Prefrontal and Motor stimulation sites. Error bars represent standard deviation across subjects (*N* = 6 for Prefrontal and *N* = 5 for Motor). Significant differences from the personalised WBM are indicated (paired t-test, Bonferroni corrected; *∗p <* 0.05, *∗ ∗ p <* 0.01, n.s. = not significant).

Finally, we evaluated whether the optimized effective connectivity itself is subject-specific by performing a cross-subject EC swap (“EC TMS”). In this condition, a subject’s resting-state WBM was combined with a TMS-refined EC matrix belonging to a different individual. In contrast to the Rest Params surrogate, swapping the optimized EC caused a drastic collapse in predictive accuracy of the full TEP spatio-temporal dynamics. As shown in Figure 4, the personalised WBM outperformed the EC TMS surrogate across both regions. In Prefrontal, personalized WBM performance (*ρ* = 0.72 ± 0.12, range [0.56, 0.89]; *R*^2^ = 0.50 ± 0.18, range [0.24, 0.78]) was significantly higher for both *ρ*(*p <* 0.01) and *R*^2^ (*p <* 0.05) compared to EC TMS (*ρ* = 0.21 ± 0.21, range [− 0.02, 0.49]; *R*^2^ = − 0.41 ± 0.65, range [− 1.62, 0.17]). Similarly, in Motor, the personalised WBM (*ρ* = 0.91 ± 0.03, range [0.88, 0.93]; *R*^2^ = 0.77 ± 0.09, range [0.65, 0.87]) showed a significant improvement over EC TMS for both metrics (*ρ*: *p <* 0.05; *R*^2^: *p <* 0.05; EC TMS: *ρ* = 0.39 ± 0.30, range [0.04, 0.67]; *R*^2^ = − 0.21 ± 0.54, range [−1.09, 0.35]). These performance losses primarily emerge at longer post-stimulation latencies, where recurrent network interactions and subject-specific feedback dynamics increasingly shape the TEP trajectory beyond the initial propagation phase.”

Together, these results demonstrate that the two key ingredients for successfully modeling the TEPs are *i*) an effective baseline, which can be obtained from the group-level resting-state profile (but not structural connectivity alone), and *ii*) the individualized and site-specific *EC*_*TMS*_ refinement procedure, which is critical to capturing the brain’s transition from spontaneous activity to sustained, complex responses observed in the late TEPs.

### Analysis of TMS-Induced Effective Connectivity Changes

The fitting procedure estimated a stimulation-site specific Effective Connectivity matrix (*EC*_TMS_) that optimally reproduced the TMS-evoked dynamics. To isolate the impact of perturbation on network topology, we analyzed the differential log-connectivity Δ log(*EC*) = log(*EC*_TMS_) − log(*EC*_Rest_). As illustrated by the dominant blue hues in the differential matrices (Figure 5A), the vast majority of directional communication channels show a marked decrease in strength during stimulation compared to the EC inferred from the resting state. However, TMS also selectively enhances a sparse set of pathways emerging from the stimulated network. This transformation effectively increases the contrast between relevant and irrelevant communication channels, creating a low-dimensional propagation backbone through which the perturbation spreads.

**Fig. 5.**
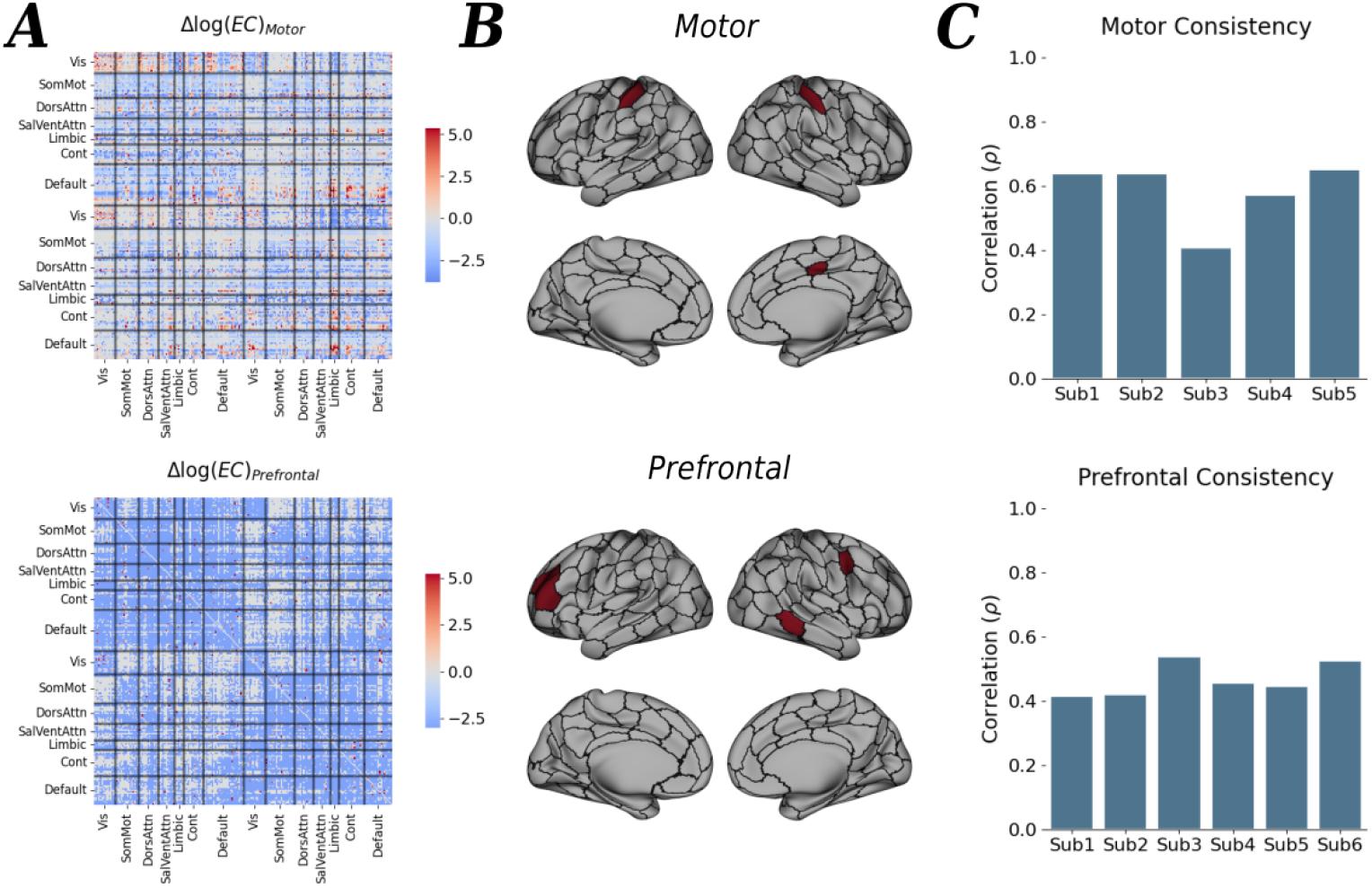
Mechanisms of TMS-induced effective connectivity. **(A)** Differential effective connectivity matrices. Full Δ log(*EC*) matrices for Motor (top) and Prefrontal (bottom) stimulation, ordered by canonical Yeo networks.**(B)** Cortical topography of major enhanced sources. Spatial projection of the nodal source scores calculated from consistently enhanced edges (surviving a 95th percentile magnitude threshold and a strict subject-consensus mask). **(C)** Subject consistency of EC modulation. Pearson correlation (*ρ*) between individual-subject differential EC matrices and the group-level average for Motor (top, *n* = 5) and Prefrontal (bottom, *n* = 6) stimulation setups.

To identify the pathways preferentially recruited during stimulation, we isolated edges showing consistent and pronounced enhancement across subjects. For each stimulation site, we constructed a consensus network containing only those connections that increased beyond a conservative threshold in the majority of subjects (≈ 60%). Within this consensus network, we calculated a nodal score defined as the total magnitude of these top-tier enhancements originating from each region (*score*_*j*_ = Σ_*i*_ Δ log(*EC*)_*ij*_).

Mapping these scores onto the cortical surface revealed a highly localized and site-specific organization (Figure 5B). Despite the widespread reduction in effective connectivity, the connections that were selectively strengthened were consistently anchored to the stimulated network. For Motor stimulation, the strongest outbound enhancements were tightly anchored to the primary motor region and supplementary motor areas. For Prefrontal stimulation, the outflow was concentrated within executive control and salience hubs. This indicates that while TMS acts as a massive global dampening filter on intrinsic communication, it simultaneously enforces a highly specific outflow originating precisely from the stimulation focus.

Finally, we evaluated how reliably this global suppression and targeted outflow pattern is preserved across individual participants by computing the correlation (*ρ*) between each single-subject’s differential profile and the group-averaged matrix (Figure 5C). The inferred EC changes displayed significant consistency across subjects. For Motor stimulation, individual patterns correlated tightly with the group trend (mean *ρ* ≈ 0.58, ranging from 0.40 to 0.65). Prefrontal stimulation exhibited an equally consistent profile across all six subjects (mean *ρ* ≈ 0.46). This high consistency underscores that the combination of widespread network suppression paired with localized structural routing is a stable and reproducible feature of the brain’s initial response to direct cortical stimulation, preceding the emergence of the subject-specific recurrent dynamics that dominate later TEP components.

### Group-Average *EC*_TMS_ Generalizes to Single-Subject Early TEP Prediction Without Refitting

The analyses above revealed a hierarchical organization of TMS-induced connectivity changes. While reproducing the full spatiotemporal evolution of the TEP requires subject-specific effective connectivity, the dominant reconfiguration induced by stimulation—characterized by widespread suppression and selective enhancement of site-specific propagation pathways—was highly consistent across individuals. This observation raises the possibility that the initial propagation of TMS-evoked activity may be governed by canonical connectivity motifs shared across subjects.

To test this hypothesis, we evaluated whether group-averaged effective connectivity patterns could accurately predict individual TMS responses without any subject-specific connectivity fitting. For each stimulation target, we constructed a group-level *EC*_TMS_ matrix by averaging the inferred connectivity across subjects and used this matrix, together with each subject’s resting-state-fitted parameters, to simulate TMS-evoked activity. Because the previous analyses suggested that shared connectivity motifs primarily shape the initial perturbational response, we restricted this evaluation to the early post-stimulus window (0–100 ms).

For each individual, early TEP responses were generated by combining the subject’s unique resting-state-fitted whole-brain parameters with the corresponding group-average *EC*_TMS_ matrix. Across the cohort, this group-average approach yielded moderate predictive capacity for the early evoked volley following motor stimulation (mean *ρ* = 0.52 ± 0.18, range [0.24, 0.69]; mean *R*^2^ = 0.29 ± 0.14, range [0.08, 0.46]), though accuracy varied substantially across individual subjects. Figure 6 illustrates this reconstruction for a representative subject, highlighting that while the overall 0–100 ms explained variance is moderate, the framework captures the earliest propagation pattern (0–20 ms) with substantially greater accuracy than the later response.

**Fig. 6.**
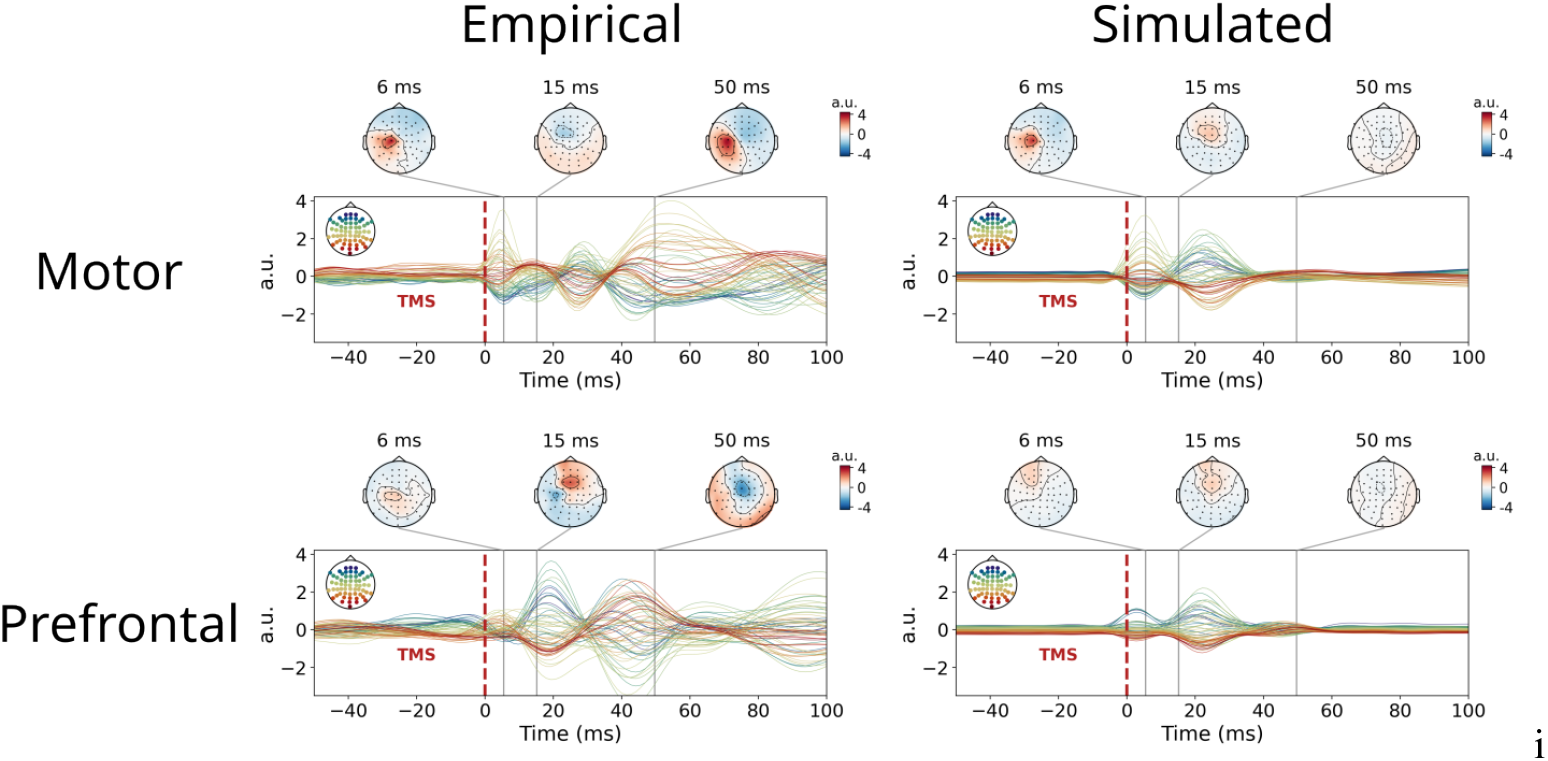
Validation of early TEP prediction using group-average effective connectivity. Comparison of empirical TEPs (left column) and model simulations utilizing a group-averaged *EC*_TMS_ matrix (right column) for the Motor (top row; *ρ* = 0.69, *R*^2^ = 0.46) and Prefrontal (bottom row; *ρ* = 0.43, *R*^2^ = 0.18) stimulation sites in a representative subject (Subject 1). The topographies highlight that the model accurately captures the localized initial cortical response (6 ms) for both targets and early propagation (15 ms) for the motor target. Conversely, the time-series illustrate the model’s systematic underestimation of late-stage amplitudes (50 ms), where complex or non-linear recurrent network dynamics typically emerge.

As observed in the full-window optimization, we tracked a similar performance gap when applying this group-average framework to Prefrontal stimulation. While the predictions were considerably weaker than those obtained for the motor target, they remained highly consistent across the cohort, showing a narrow distribution of performance (*ρ* = 0.48 ± 0.11, range [0.27, 0.62]; *R*^2^ = 0.21 ± 0.07, range [0.07, 0.27]). In line with our previous observations, this reduction in reconstruction accuracy suggests that prefrontal responses may depend more strongly on subject-specific or higher-order recurrent dynamics, even at relatively early latencies.

In contrast, replacing the group-average *EC*_TMS_ with the group-average structural connectivity (SC) led to a pronounced drop in predictive performance (mean *ρ* = 0.31 ± 0.17, range [0.01, 0.51]; mean *R*^2^ = 0.06 ± 0.13, range [−0.01, 0.21]), failing to reproduce the observed propagation patterns. This contrast demonstrates that the stimulation-induced reconfiguration captured by *EC*_TMS_, including both widespread suppression and selective pathway enhancement, contains information that is essential for reproducing early evoked responses and is not captured by anatomical connectivity alone.

To confirm the spatial specificity of these group-level configurations, we performed a cross-site generalization test. Using the prefrontal group-averaged EC to predict motor responses yielded negligible accuracy (mean *R*^2^ = 0.05 ± 0.05, range [0.01, 0.12]), while the reverse configuration failed completely (all cross-site *R*^2^ *<* 0). These results demonstrate that the group-level *EC*_TMS_ matrices encode site-specific propagation motifs rather than a generic response to cortical perturbation.

Together, these findings reveal a hierarchical organization of the evoked response: while highly individualized effective connectivity is required to explain the later, complex components of the TEP, the immediate cortical response to localized perturbation is largely constrained by a site-specific propagation profile that remains qualitatively consistent across individuals. Thus, a substantial fraction of the initial 100 ms of the TEP can be approximated using a group-averaged effective connectivity, indicating that early cortical responses are governed by a shared propagation scaffold, whereas later components increasingly reflect subject-specific recurrent dynamics.

## Discussion

Our primary contribution is the establishment of a generative framework that mechanistically links the brain’s response to perturbation to its intrinsic resting-state dynamics. Methodologically, we provide a closed form cross-spectral solution for a connectome-coupled Hopf model with delays and a biophysical projection to sensors. This allows for the direct fitting of the resting state EEG covariance matrix without the need for computationally expensive time domain simulations. We show that the resting state is best characterized by supercritical local dynamics (*a*_*i*_ *>* 0), where alpha-range oscillations emerge from active limit-cycles rather than filtered noise. This oscillatory background, derived entirely from spontaneous data, defines the landscape upon which external perturbations propagate. The inability of the model to reproduce phase-scrambled EEG data, in which temporal phase relationships were destroyed while preserving spectral content, shows that the model success in fitting resting state covariance matrix does not arise from unrestricted fitting flexibility. Rather, the model appears to capture statistical and dynamical features that are specific to physiological brain activity, increasing confidence that the inferred parameters reflect meaningful aspects of the underlying neural organization.

We further demonstrate that the intrinsic spectral fingerprint of the resting state provides a dynamical scaffold to predict the spatiotemporal evolution of TMS–evoked potentials. This relationship is not merely correlational. Unlike previous efforts that modeled TEPs in isolation, our approach treats resting-state dynamics as a biologically grounded prior that constrains the estimation of perturbation-specific effective connectivity. In fact, a key finding of this work is the crucial role of the resting state prior in regularizing the inference of the effective connectivity (EC). Our results indicate that while connectivity gains are the primary drivers of site-specific TEP propagation, they cannot be stably inferred in the absence of resting-state constraints. Moreover, by constraining the stimulation-specific EC to remain close to the resting-state backbone, the model is biased toward physiologically plausible solutions rather than overfitted mathematical configurations. This constraint is critical because unconstrained optimization can achieve high predictive accuracy even when local dynamical parameters are randomized or biologically implausible (see Figure S15). Thus, the model exposes a fundamental tension between flexibility and interpretability: without biologically grounded priors, accurate fits may correspond to mathematically valid but physiologically meaningless network solutions.

Our two-step protocol, in which intrinsic local dynamics and EC are first calibrated at rest and EC is subsequently inferred using a minimal subset of TEP trials, allowed us to infer personalized, stimulation-site-specific EC patterns: in both cases we observed a widespread reduction of effective interactions, together with the selective strengthening of a restricted set of pathways anchored to the stimulated network. This pattern is consistent with the idea that TMS transiently reorganizes large-scale communication, favoring the propagation of activity along network-specific routes while reducing the contribution of ongoing background interactions. This interpretation is broadly consistent with recent studies showing that TMS responses depend strongly on the large-scale brain activity before the stimulation and the stimulus network state, rather than solely on static local cortical properties [16].

From this perspective, the widespread suppression of connectivity may reflect a transient reduction in distributed background communication, whereas stimulation transiently reshapes this architecture in a site-specific manner, producing reproducible propagation motifs that can be observed across individuals in TEP early response. These motifs partly reflect a canonical architecture that generalizes across individuals: group-averaged, site-specific EC templates provide a meaningful approximation of short-latency TMS responses at the single-subject level, without the need for individualized connectivity refitting. This finding suggests that the earliest phase of the TEP is largely determined by propagation properties that are intrinsic to the stimulated network and conserved across subjects. However, the moderate predictive power of the group-level model and the substantial variability observed across individuals reveal a clear limit to this generalization. While the initial response can be approximated using shared propagation motifs, later TEP components increasingly depend on subject-specific recurrent interactions and state-dependent network dynamics. Importantly, the large performance gap between effective connectivity-based predictions and those obtained using structural connectivity alone demonstrates that anatomical coupling provides only a partial description of perturbational spread. Instead, the propagation of TMS-evoked activity is governed by directed interactions that emerge from the underlying dynamical state of the network. Together, these findings support a hierarchical view of cortical reactivity in which conserved, stimulation-site-specific motifs shape the initial propagation of activity, whereas individualized recurrent dynamics progressively dominate the subsequent evolution of the response.

Because our model is generative and only requires a connectivity calibration based on a few stimulation trials, it offers a principled route to individualized stimulation planning. In clinical settings, a short resting EEG recording (e.g., 5–10 minutes) could be used to parameterize the subject’s dynamical prior, while a minimal number of TMS pulses (e.g., 10–15 trials, corresponding to our 10% training subset) could refine the target specific couplings. This framework allows for the *in silico* testing of different stimulation sites and protocols, providing a powerful tool for the design of individualized, model driven neurostimulation interventions.

Some limitations of the current study should be explicitly discussed. First, the small cohort size (N=11 split between stimulation sites) may limit the generalizability of the group-level effective connectivity templates. Future studies utilizing larger cohorts, possibly including additional stimulation sites, will confirm whether the site-specific propagation patterns remain stable across broader populations. Second, we observed a marked asymmetry in model performance between stimulation sites, with prefrontal targets yielding lower predictive accuracy compared to motor ones. This performance gap suggests that prefrontal stimulation may engage more complex, state-dependent, or non-linear recurrent dynamics that are less captured by static effective connectivity. Third, it is important to clarify the boundaries of the model’s predictive capacity: while the resting-state provides a dynamical scaffold, estimating the TMS-induced effective connectivity still requires a minimal subset of empirical TMS data, rather than being purely derivable from resting-state activity alone. Finally, we highlight that in this study a group average anatomical connectivity is used given the lack of individual diffusion tensor imaging (DTI) scans. While our model successfully bridges rest and reactivity, several avenues for extension remain. Currently, the effective connectivity is estimated as a static modulation of the structural connectome; future developments could incorporate time-varying EC to capture the non-linear evolution of late TEP components. Additionally, exploring how the resting state scaffold changes across different brain states (e.g., sleep, anesthesia, or cognitive tasks) will be essential for understanding the state dependency of TMS responses. Finally, integrating multi-modal priors such as receptor density maps or individual microstructural data could further refine the biophysical accuracy of the local Hopf dynamics.

## Methods

### Participants and EEG acquisition

We analyzed resting-state EEG and TMS–EEG from twelve healthy adults acquired with a 60-channel TMS-compatible system (0.1–350 Hz hardware bandpass) [CITE]. One subject from the motor-cortex stimulation cohort was subsequently excluded from final analysis due to an excessive number of contaminated channels during baseline recording, resulting in a final analyzed cohort of 11 subjects. Preprocessing included high-pass filtering at 1 Hz, channel interpolation, 2 s epoching, downsampling, artifact rejection, ICA-based component removal, and low-pass filtering at 80 Hz.

### Parcellation, connectivity, and delays

We parcellated cortex with the Schaefer 7Networks, 200-node atlas (2 mm). Structural connectivity (SC) came from HCP-derived group-averaged tractography; inter-node delays were distance-based (*m*_*jk*_ = *d*_*jk*_*/v*) with fitted axonal conduction velocity *v*.

### Neural mass model

Local node dynamics are governed by the Hopf normal form, a canonical model for oscillatory systems also known as the Stuart-Landau oscillator [33]. This model captures the fundamental transition from a stable state to self-sustained oscillations as the bifurcation parameter *a* is varied. The dynamics for a single node are given by:

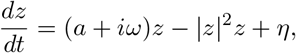

where *z* = *x* + *iy* is the complex state variable, *ω* is the intrinsic angular frequency, and *η* is additive white noise. In a network context, the dynamics of node *j* become:

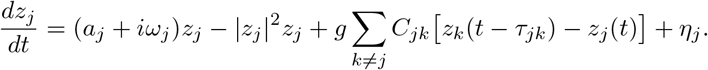

To account for external stimulation, such as Transcranial Magnetic Stimulation (TMS), we introduce a perturbation term *p*_*j*_(*t*) into the system. The TMS pulse is modeled as an additive input to the real part of the trajectory, representing the induced current in the source space. The equation for the real component *x*_*j*_ of the node *j* is thus given by:

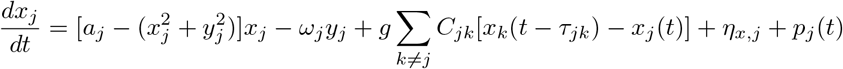

where *p*_*j*_(*t*) is the source-space reconstruction of the TMS pulse.

To analyze the system’s linearized response, we compute the block Jacobian *A* and frequency operator *U*(*ν*), which are used to derive the network’s spectral properties *ψ*(*ν*) = *U*(*ν*)*QU* ^†^(*ν*) and the covariance matrix 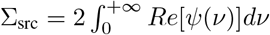.

### Cortical heterogeneity

Regional heterogeneity was incorporated by scaling the bifurcation parameter *a*_*j*_. T1w/T2w values from the Human Connectome Project were projected onto the Schaefer 200 atlas using the neuromaps toolbox and normalized via an error function to produce standardized scores *h*_*j*_ ∈ [0, 1]. The parcel-specific parameter was then defined as:

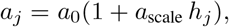

where *a*_0_ is the baseline value and *a*_scale_ is a scaling coefficient estimated during model fitting. This approach captures known hierarchical gradients in cortical microstructure and excitability, consistent with recent empirical evidence showing that high-order cortical regions exhibit greater excitability and recurrent feedback than low-order ones [34] (see Supplementary Note 7 and Supplementary Figure S16).

### EEG forward model and stimulation input

We projected source activity to sensors via subject-specific leadfields *G*, obtained using each subject’s resting state EEG data in combination with the standard template MRI subject (fsaverage) through MNE’s *EEG forward operator with a template MRI* pipeline (see Supplementary Note 8). The macroscopic neural activity from the Hopf model was taken as the real part of its complex state variable, *x*_*i*_(*t*) = Re[*z*_*i*_(*t*)], which was then projected to sensor space as:

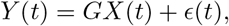

where *X*(*t*) = [*x*_1_(*t*), …, *x*_*N*_ (*t*)]^*T*^ and *ϵ*(*t*) is measurement noise. The resulting sensor-level covariance is given by Σ_sens_ = *G*Σ_src_*G*^⊤^ + Σ_noise_. TMS inputs were constructed by thresholded electric field maps (normalized to peak) for each target using SimNIBS [32], yielding parcel-wise stimulation weights for motor and prefrontal targets (see Supplementary Note 9 and Supplementary Figure S17).

### Resting-state fitting objective and optimization

The model was first fitted to each subject’s resting state EEG to characterize the intrinsic dynamical scaffold. We minimized the Frobenius norm between the empirical and model predicted sensor-space log-transformed covariance matrices, leveraging our closed form analytical solution for the cross spectral density. This log-transformation was employed to linearize the multiplicative differences in signal variance across sensors, ensuring that the optimization captures the global network topology rather than being disproportionately biased by high-power channels. The parameters estimated at this stage included the node specific bifurcation parameters (*a*_*i*_), intrinsic frequencies 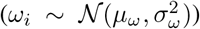, the global coupling scaling (*g*), the standard deviation of the input noise (*σ*_*in*_), and the homogeneous conduction velocity (*v*_*d*_). Notably, the fitting was performed on the unaltered resting state data and compared against a null model derived from phase scrambled EEG to verify that the optimized parameters reflected genuine spatial and temporal correlations rather than just power spectral features.

### Effective connectivity (EC) and TEP refitting

While structural connectivity (SC) provides the anatomical backbone for signal propagation, it often under explains the specific spatiotemporal spread of TEPs. Therefore, once the resting state parameters were fixed, we performed a secondary refitting of the connectivity weights to a small subset (10%) of the TMS-EEG trials for each target. We augmented the SC with a row-normalized, log-weighted modulation and a Laplacian balance term to define the effective connectivity (EC). Crucially, during this step, all parameters inherited from the resting state fit (*a*_*i*_, *μ*_*ω*_, *σ*_*ω*_, *g, σ*_*in*_, *v*_*d*_) remained constant. The EC was optimized using a Gaussian prior centered on the resting state connectivity weights, strictly constraining the model to a biologically plausible neighborhood. This ensures that the resulting TEP prediction is a product of the intrinsic resting state dynamics modulated by site-specific connectivity adjustments, rather than an unconstrained mathematical fit.

### Evaluation metrics

We quantified variance explained (*R*^2^) of sensor covariance, median scaled residuals, temporal latency matches for TEP components, spatial correlations of band-limited power maps, and ERSP similarity across subjects and targets.

### TMS Stimulation Patterns

TMS stimulation patterns are incorporated by computing the induced electric field distribution using SimNIBS 4.1[32], [35]) in MNI152 standard space.

## Data availability

Data supporting the findings of this study are available from the corresponding author upon reasonable request.

## Code availability

The source code for the modeling and analysis presented in this study is openly available on GitHub at https://github.com/veronean/Modelling-Response-to-Brain-Stimulation-from-Resting-State-Activity. The specific version of the code used to produce the results in this paper is archived and persistently accessible via Zenodo at https://doi.org/10.5281/zenodo.19128186.

## Acknowledgements

xxx

## Author contributions

**Andrea Veronese:** Conceptualization, Methodology, Software (computational implementation), Formal analysis, Data Curation (preprocessing), Writing – Original Draft, Writing – Review & Editing. **Davide Momi:** Methodology, Software (foundational code), Resources (SimNIBS reconstructions), Writing - Review & Editing. **Simone Sarasso:** Resources (empirical data acquisition), Data Curation (guidance on preprocessing), Writing - Review & Editing. **Maurizio Corbetta** Writing - Review & Editing. **Michele Allegra:** Conceptualization, Supervision, Writing – Review & Editing. **Samir Suweis:** Conceptualization, Supervision, Writing – Review & Editing.

